# Disruptions in Primary Visual Cortex Physiology and Function in a Mouse Model of Timothy Syndrome

**DOI:** 10.1101/2024.12.20.629743

**Authors:** Rosie Craddock, Cezar M. Tigaret, Frank Sengpiel

## Abstract

Timothy syndrome (TS) is a rare genetic disorder caused by mutations in the *CACNA1C* gene which encodes the L-type calcium channel α-1 CaV1.2 subunit. While it is expressed throughout the body the most serious symptoms are cardiac and neurological. Classical TS1 and TS2 mutations cause prolonged action potentials (APs) in cardiomyocytes and in induced neurons derived from pluripotent stem cells taken from TS patients, but effects of TS mutations on neuronal function in vivo are not fully understood. TS is frequently associated with autistic traits, which in turn have been linked to altered sensory processing. Using the TS2-neo mouse model we analysed effects of the TS2 mutation on the visual system. We observed a widening of APs of pyramidal cells in ex vivo patch-clamp recordings and an increase in the density of parvalbumin positive (PV+) cells in the primary visual cortex. Neurons recorded extracellularly in vivo were less likely to respond to visual stimuli of low spatial frequency, but more likely to respond to visual stimuli of mid-to-high spatial frequency, compared to WT mice. These results point to a basic processing abnormality in the visual cortex of TS2-neo mice.

## Introduction

Timothy syndrome (TS) is a rare, multisystemic, genetic condition that is associated with a combination of cardiac (long QT interval and arrythmias), endocrine (pancreatic, adrenal and thyroid dysfunction) and neuropsychiatric manifestations and a somatic morphological phenotype (cardiovascular malformations and syndactyly). Neurological symptoms of TS include autism spectrum disorder (ASD), epilepsy and developmental delay (Splawski et al. 2004; Splawski et al. 2005; Walsh et al. 2018). TS is caused by mutations in the *CACNA1C* gene which is associated with risk across a number of neuropsychiatric conditions (Liu et al. 2011; Lu et al. 2012). *CACNA1C* encodes the α-1 CaV1.2 subunit of L-type voltage-gated calcium channels (L-VGCC) which are expressed in a variety of tisues (Uhlén et al. 2015).

Subtypes of TS are typically categorised by causative mutation in the *CACNA1C* gene. Two missense G406R point mutations in exons 8A and 8 cause, respectively, TS type 1 (TS1) and TS type 2 (TS2) (Splawski *et al*. 2004) (Splawski *et al*. 2005) (Bauer et al. 2021) L-type Ca2+ channels containing α subunit with TS mutations (TS-VGCCs) have a reduced voltage-dependent inactivation (VDI; (Barrett and Tsien 2008)). Consequently, TS-VGCCs remain open for a prolonged period upon channel activation, generating prolonged depolarising calcium influx into the cell (Splawski *et al*. 2004; Splawski *et al*. 2005; Paşca et al. 2011; Yazawa et al. 2011). In cardiomyocytes, the functional properties of TS-VGCCs result in an increased duration of cardiac action potentials (APs), thought to underlie cardiac symptoms in TS including long QT, arrhythmia and cardiac arrest (Splawski *et al*. 2004; Splawski *et al*. 2005; Dick et al. 2016). In contrast, the mechanistic links between TS-VGCC and the neurological phenotype in TS are not yet understood.

In neurons, CaV1.2 L-VGCCs are expressed predominantly somatodendritically and are activated by membrane depolarization during APs to generate Ca2+ influx important for synaptic function, shaping of AP firing, and regulation of gene expression (Zamponi et al. 2015). In addition, L-VGCCs have been critically implicated in neurodevelopmental neuronal migration [(Jiang and Swann 2005); (Kamijo et al. 2018)]. Observations in neurons differentiated from TS patient-derived induced pluripotent stem cells (iPSCs) indicates that the neurobiological consequences of TS mutation are electrophysiological (prolongation of AP and increased residual cellular Ca2+ (Paşca *et al*. 2011)) and neurodevelopmental (aberrant interneuron migration (Birey et al. 2017)). This suggests a neurodevelopmental pathology in common with other neurodevelopmental conditions (Hussman 2001; Paterno et al. 2021; Righes Marafiga et al. 2021). However, the impact of TS-VGCCs on neuronal and circuit functional properties in the adult brain are less clear.

Moreover, abnormalities in inhibitory interneuron migration have been demonstrated in cultured human forebrain spheroids from TS1 patients (Birey *et al*. 2017). The development of parvalbumin-positive (PV+) cells, the most numerous type of inhibitory interneuron, is regulated by L-type voltage-gated calcium channels, including CaV1.2 (Jiang and Swann 2005). Therefore, functional changes in the channel resulting from TS mutations could impact PV+ cell development and migration.

Abnormalities in visual perception and in vision-related behaviours have been reported in some TS patients. These sensory processing abnormalities are known to be associated with ASD which is also frequently exhibited in TS patients. Both gaze aversion and the limited interest in faces displayed by ASD patients has been attributed to abnormal processing of facial visual stimuli (Dalton et al. 2005; Coulter 2009; Zhou et al. 2023). It has been proposed that the lack of visual attention to facial stimuli, and reduced ability to perceive and recognise faces in ASD patients occurs as a result of differences in basic visual processing in the ASD brain (Coulter 2009; Vlamings et al. 2010; Zhou *et al*. 2023), including an enhanced contrast sensitivity for simple visual gratings stimuli of high spatial frequency (Kéïta et al. 2014) and increased ability to see the fine detail in visual images, as opposed to the big picture (Shah and Frith 1983; Robertson and Baron-Cohen 2017; Zhou *et al*. 2023). However, the mechanistic roles of TS VGCCs in this sensory phenotype are not well understood (Marco et al. 2011).

In order to understand the consequences of TS VGCCs at the neuronal and systems levels on sensory processing in the adult brain, we used a mouse model expressing the TS2 mutation (TS2-neo mouse) exhibiting TS-relevant behavioural and neurodevelopmental abnormalities (Bader et al. 2011; Bett et al. 2012; Rendall et al. 2017; Horigane et al. 2020). TS2-neo mice have a superior ability to perceive complex auditory stimuli (Rendall *et al*. 2017), without gross perceptual disturbances (Bett *et al*. 2012), consistent with the perceptual enhancements reported in ASD patients (Bertone et al. 2005; Mottron et al. 2006). However, the links between TS2 mutations and visual processing abnormalities are less clear.

Here, we describe a series of experiments using TS2-neo mice aimed to explore neurobiological consequences of the TS VGCCs on visual processing in the primary visual cortex (V1), AP firing properties of V1 pyramidal neurons, and the density of PV+ interneurons in the visual cortex.

## Materials and methods

### Animals

All experimental procedures were carried out in accordance with the UK Animals (Scientific Procedures) Act 1986 and European Commission Directive 2010/63/EU. Mice heterozygous for the Timothy Syndrome type TS2-neo mutation (background C57BL/6J) were purchased from Jackson labs (Jax #019547) and bred with C57BL/6J mice also purchased from Jackson labs (Jax #000664). Mice generated from this breeding colony were used in all experiments described here. Same-sex littermates were housed together under normal light conditions (12-hours light and 12-hours dark) and were given standard food and water ad-libitum. Environmental enrichment was provided to the mice. Experimenters were kept blind to genotype throughout the study until completion of final data analyses in each instance. Animal treatments and usage specific to each experiment are detailed at the beginning of the following subsections.

### Electrophysiology

8 mice heterozygous for the TS2-neo mutation and 5 sibling WT mice of both sexes were used in this experiment. Mice aged 4-10 weeks old were sacrificed by an overdose of isoflurane before decapitation and rapid removal of the brain under ice cold choline-chloride slicing solution (110mM choline chloride, 25mM D-glucose, 15mM NaHCO_3_, 2.5mM KCl, 1.25mM NaH_2_PO_4_, 11.6mM L-ascorbic acid, 2.1mM sodium pyruvate, 0.5mM CaCl_2_, 7mM MgCl_2_, 310mOsm). Brains were coronally sectioned to a thickness of 400µm using a vibratome (CI.7000SMZ-2, Campden instruments). Sections containing V1 were collected and incubated for 30 minutes in carboxygenated (95% O_2_, 5% CO_2_) artificial cerebrospinal fluid (ACSF; 11.9mM NaCl, 10mM D-glucose, 2.6mM NaHCO_3_, 2.5mM KCl, 1mM NaH_2_PO_4_, 2.5mM CaCl_2_, 1.3mM MgCl_2_) at 34°C in a brain slice holding chamber (BSK1, Brain Slice Keeper). After incubation, the brain slice holding chamber was left at room temperature for at least 1 hour before use.

Slices were mounted onto poly-L-lysine coated glass coverslips which were then placed into a submerged recording chamber (Slice Minichamber II, Luigs and Neumann) which was perfused with carboxygenated ACSF at 32°C. Pyramidal cells in V1 were visualised using an infra-red differential interference contrast imaging system (BX-51 WT, Olympus). Putative pyramidal cells were targeted based on somatic shape, size and location in layer 2-5 of V1. Cell type was later confirmed by biocytin-streptavidin histology and confocal imaging using a confocal microscope (Zeiss Airyscan LSM880; methods based on Swietek *et al*., 2016). A maximum of 2 cells were recorded from each animal.

Micropipettes filled with internal solution (117mM potassium methanesulfonate, 8mM NaCl, 10mM 4-(2-hydroxyethyl)-1-piperazineethanesulfonic acid, 4mM MgATP, 0.3mM NaGTP, and 0.2mM Ethylene Glycol-bis (β-aminoethyl ether)-N,N,N′,N′-Tetra acetic Acid, pH7.4, 280mOs) were used to attain whole-cell patch clamp. The pipette resistance was measured at 4-5MΩ in each instance. The liquid junction potential was corrected for using Multiclamp 700B software. Patched cells were held at −70mV for 1 minute before measuring membrane potential responses to a series of incremental current step injections between −50 and 650pA. Each current step injection was 200ms duration followed by a 2s 0pA injection, the increment between steps was 50pA. The experimental protocol was designed and implemented using Clampex 11 software. Membrane potential data were lowpass filtered to 6kHz, sampled at 50kHz, and digitized using an Axon Digidata 1550 Data Acquisition System and Clampex 11 software.

Analysis of electrophysiological recordings was performed with custom software written using Python 3.10 and the pyABF package (Harden 2022). The recorded membrane potential trace resulting from the hyperpolarising −50pA current step injection was used to measure passive membrane properties of the cell (as per Tamagnini *et al.,* 2015). A more detailed description of how passive membrane potential properties were measured is provided in the Supplementary Methods and in Supplementary Figure 1. Measurements of single AP properties were taken from the first AP which was fired during the experimental protocol. Membrane potential threshold was measured as the membrane potential at which the rate of change of membrane potential reached 15.2mV/ms (Paşca *et al*. 2011). The membrane potential half-way between the threshold potential and the AP amplitude was designated the AP half-height. The duration of time for which the membrane potential exceeds this AP half-height for the first measured AP was taken as the AP half-width. The total AP rise and fall times were taken as the time between threshold and amplitude, and amplitude to return to threshold potential, respectively.

Data analysis was completed in R. The distribution of all properties within the dataset for V1 cells were tested for normality by Shapiro-Wilk testing. Membrane capacitance, time constant, threshold potential, AP amplitude, AP half-width, maximum rates of rise and fall of the AP, the total times for AP rise and fall were all found to be normally distributed by Shapiro-Wilk testing (P>0.05), while the distribution of values of membrane input resistance in the dataset were found to be non-normally distributed by Shapiro-Wilk testing (P=0.038, W=0.88). Genotype differences in all normally distributed variables were tested by t-tests while the genotype difference in membrane input resistance was tested by Wilcoxon signed rank test.

### In vivo two-photon calcium imaging of V1

Data for this study were obtained from 5 TS2-neo and 4 WT mice of either sex. Mice underwent surgery at 10-12 weeks of age. Surgical procedures were based on those described previously by our lab (Powell et al. 2020; Craddock et al. 2023). Mice were given prophylactic antibiotics and anti-inflammatory medication for one week prior to surgery. Mice were anaesthetised using isoflurane and secured in a stereotaxic frame (David Kopf Instruments, Tujunga, CA, USA). The scalp and periosteum were removed from the dorsal surface of the skull. A custom headplate was attached to the cranium using dental cement (Super Bond, C&B). The aperture of the headplate was cantered over the right hemisphere V1 of the mouse brain, 2.3mm lateral and 3.1 posterior to bregma. A 3mm circular craniotomy was made in this position. Mice were injected with 150µl of AAV1.Syn.GCaMP6f.WRPE.SV40 viral vector (Penn Vector Core, 1:10 dilution of 3.45×1013gc/ml) in V1b (3.1mm lateral and 3.1mm posterior to bregma) such that the fluorescent calcium indicator, GCaMP6f, was expressed in neurons of V1b of the mouse brain. The craniotomy was closed using 2x 3mm circular glass inserts bonded together, overlaid with a 5mm circular insert (Biochrom Ltd., 3mm: product code 640720(CS-3R), 5mm: product code 640700) which was attached to the cranium-fixed headplate. Glass inserts were bonded together using optical adhesive (Norland Products; catalogue no. 7106). The window was sealed with dental cement and the mice left to recover under surveillance. Mice were returned to their home cage with sibling mice and left to recover for 1 week. The mice received post-operative checkups for the first 3 days of this recovery period and were administered post-operative anti-inflammatories and antibiotics.

Mice underwent a 1-week period during which they were habituated to handling and head fixation using methods similar to those described by (Aguillon-Rodriguez et al. 2021). Mice were placed on top of a custom-built cylindrical treadmill and were head-fixed to a resonant scanning two-photon microscope (Thorlabs, B scope) by the mouse’s cranium-mounted head plate. Mice were positioned to face a calibrated LCD PC screen (Iiyama, B2080HS; width × height 26 × 47 cm^2^) positioned in the mouse’s binocular field of view, 20cm away from the mouse.

In vivo two-photon imaging was then performed on awake mice receiving visual stimulation via the screen positioned in their binocular field of view. Imaging involved a resonant scanning two photon microscope equipped with a 16×0.8NA water-immersion objective (Nikon). GCaMP6f in neurons of L2/3 was excited at 920nm using a Ti:sapphire laser (Coherent, Chameleon) using a maximum power of 50mW (as measured at the sample). Data were acquired using ThorImage software at a raw frame rate of 56Hz. We averaged over 6 frames, giving an effective frame rate of 9.6Hz. Imaging timestamps were collected using MATLAB 2022a and ThorSync software and National Instruments data acquisition cards. Recordings were made from a 1250µmx1250µm (256×256 pixel) field of view, at a depth of 150-250µm from the pial surface, corresponding to cortical L2/3 of V1b. Each mouse was recorded from up to 3 times, with up to 2 recordings being taken from a single mouse in one day. Each of the 3 recordings were taken from a different field of view within V1b.

Visual stimuli were designed and generated using custom-made codes written MATLAB (2022a) and in C# alongside the psychophysics toolbox (Brainard 1997). Stimuli were presented on the calibrated LCD PC screen described previously. Codes used to run the experiment are available on GitHub. ThorImage and ThorSync software are open source and are available on request from Thorlabs. Visual stimuli were unidirectionally drifting (to the mouse’s right) vertical sinusoidal gratings shown across the full monitor screen. The drifting gratings had a temporal frequency of 1.25Hz and varied in spatial frequency (SF) and in contrast. Stimuli of 7 different SFs were shown to each mouse: 0.014 cycles per degree (cpd), 0.031cpd, 0.064cpd, 0.128cpd, 0.236cpd, 0.383cpd. Stimuli of each SF tested were shown at 100%, 50%, 25%, 12.5%, 6.7% and 3.4% contrast such that we could find the minimum contrast at which the mouse’s V1b pyramidal neurons responded to visual stimuli of each SF tested. The contrasts and SFs used in this experiment were based on those used in similar experiments previously (Prusky and Douglas 2004; Umino et al. 2018; Cheng et al. 2020). 42 different visual stimuli were used in the experiment-one for each SF at each contrast. These were shown in a pseudo random order to the mouse, this was repeated 5 times in a single experimental recording. Single visual stimuli were shown for 3s, followed by a 3s baseline period where a grey screen of equivalent luminescence was shown to the mouse.

Images were registered and cells detected using Suite2P software (Pachitariu et al. 2017). Further details are described in the Supplementary Methods.

Further analyses were conducted in R. The proportion of all cells which were found to be visually responsive was compared by genotype using a Wilson’s test of equal proportions (Wilson, 1927). Visually responsive cells were sorted into categories of contrast sensitivity for stimuli of each SF, where contrast sensitivity is defined as 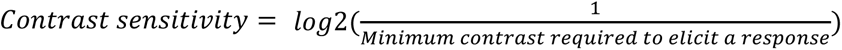. Cells which were non-responsive to visual stimuli of a specific SF were placed into a contrast sensitivity category “NR”; for details, see Supplementary Methods. Post-hoc tests of effect of genotype on contrast sensitivity at each SF was completed by use of emmeans and multcomp packages in R (Hothorn et al. 2023) (Searle et al. 1980).

### Histology, immunofluorescence techniques and imaging

Tissues from 7 mice heterozygous for the TS2-neo mutation and from 4 WT mice were used to obtain PV+ cell counts from V1 in this experiment. Mice were of either sex and were 20 weeks of age at the time of sacrifice and tissue extraction. Mice were anaesthetised using isoflurane and sacrificed using an overdose of intraperitoneal Euthatal. Mice were then transcardially perfused with 4% paraformaldehyde (PFA) in phosphate buffered saline solution (PBS 1x). Whole brains were extracted and stored in 4% PFA for 48hrs before being washed and stored in PBS 1x for 5 days. Brains were sectioned coronally to 50µm using a Leica LS1000 vibratome. 1 in 4 sections containing V1 were collected and stored in PBS 1x for up to 56hrs. All washes described in this protocol were 10 minutes long and were completed at room temperature. Sections were washed 3 times in 1x PBS, and then washed a further 4 times in 1x PBS containing 0.2% triton. Sections were incubated in a blocking solution (3% normal goat serum in 1x PBS with 0.2% triton) for 1hr at room temperature. Sections were then incubated with primary antibody solution (1:1000 Rabbit anti-PV [Swant, PV27], 3% normal goat serum in 1x PBS with 0.2% triton) for 48hrs at 4°C. Sections were washed 3 times in 1x PBS with 0.2% triton. Sections were incubated in secondary antibody solution (1:500 Goat anti-rabbit AlexaFluor488 [Invitrogen, A-11008] in 1x PBS with 0.2% triton) for 2hrs at room temperature. Sections were washed 3 times in 1x PBS with 0.2% triton and then stained with Hoechst 33342 solution to provide a control nuclear stain. Sections were washed 4 times in PBS 1x with 0.2% triton before being mounted onto SuperFrost Plus microscope slides, cover slipped using Fluoromount aqueous mounting medium. Coverslip edges were sealed using clear nail polish.

Images were obtained using an Olympus VS200 ASW slide scanner at x4 magnification. Images were taken at 1 depth per section only. This depth of focus for imaging was selected automatically by the autofocus feature of the Olympus VS2000 ASW microscope and associated OlyVIA software. PV+ cell counts were collected using automated methods in FIJI software. Cell counts were taken respective to the area of V1 within the imaged section to obtain a measurement of PV+ cell density per mm^2^. At least 8 sections were used to obtain V1 PV+ cell density measurements from each animal (range: 8-26 sections per animal).For the statistical analysis of PV+ cell density data, see Supplementary Methods.

### Correction for multiple testing and reporting results

There were 3 primary hypotheses in this study, that TS2-neo heterozygosity would: 1) prolong the AP in pyramidal cells of V1; 2) alter contrast sensitivity recorded in mice; 3) alter PV+ cell density within V1. The 3 P values related to the 3 primary hypotheses were corrected using the stringent Bonferroni correction method.

12 tests related to secondary hypotheses of this study were completed in this series of experiments. P values from these tests were corrected for 12 multiple tests by use of the Benjamini-Hochberg method.

All test values, P values, estimated effect sizes, and summary values are given to 2 significant figures (sf) in this paper. Where summary values are given the mean value is provided alongside the standard deviation in that value, the number of observations is given as n and the number of animals is given as N.

## Results

### Prolonged AP in V1 Pyramidal Cells of the TS2-neo mice

To understand the electrophysiological consequences of TS VGCCs in the visual cortex, we measured passive membrane and AP firing properties of V1 pyramidal neurons in ex vivo slices, in whole-cell patch-clamp recordings.

Passive membrane properties including membrane input resistance (Rin), membrane capacitance (cap) and membrane time constant (τ) were not found to be impacted by genotype (Rin [Wilcoxon signed rank test]: W=28, P=0.72, corrected P=0.81; cap [t-test]: t=-0.58, P=0.58, corrected P=0.81; τ [t-test]: t=-0.99, P=0.34, corrected P=0.81). AP threshold and amplitude also did not vary significantly by genotype (AP threshold [t-test]: t=-0.25, P=0.81, corrected P=0.81; AP amplitude [t-test]: t=0.46, P=0.65, corrected P=0.81).

AP duration, measured by AP half-width, was found to be significantly longer in V1 pyramidal cells from TS2-neo mice as compared to those from WT mice (t-test: t=-2.3, P=0.016, corrected P=0.048). AP duration was on average 29% longer for V1 pyramidal cells from TS2-neo as compared to those from WT mice based on comparison of the raw data (AP half-width: TS2-neo = 1.0±0.21ms [n=11, N=8]; WT= 0.80±0.24ms [n= 10 cells, N= 5]). An example AP waveform from a V1 pyramidal cell of a TS2-neo mouse compared to that from a WT mouse is shown in Figure 1. AP half-widths measured from V1 pyramidal cells of TS2-neo and WT mice are shown in Figure 2.

**Figure 1:**
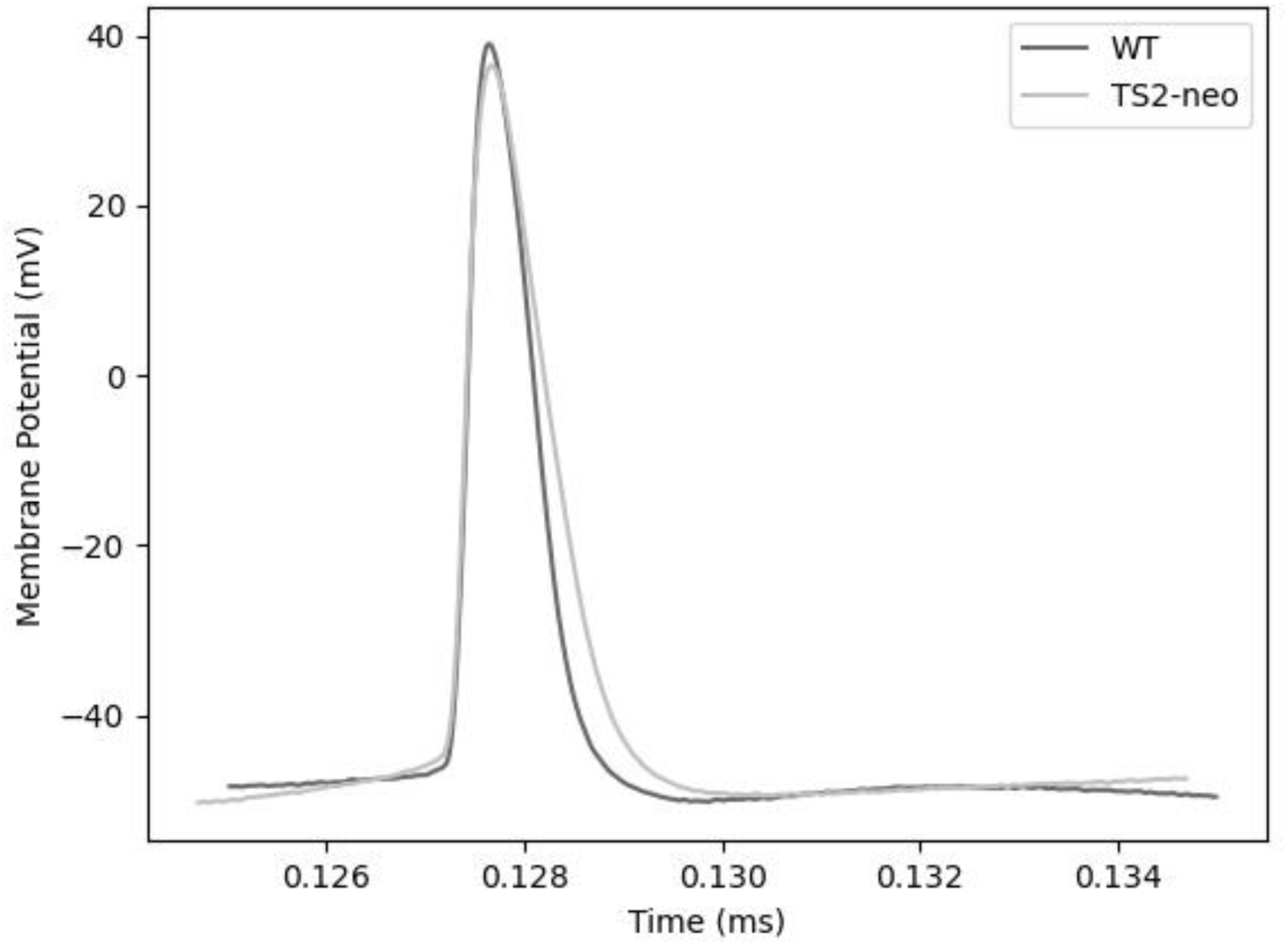
Representative AP waveforms from a V1 pyramidal cell of a TS2-neo mouse (grey line) and a WT mouse (black line). Note the increased AP width in the TS2-neo mouse. Time of current injection was at 50 ms (not shown).

**Figure 2:**
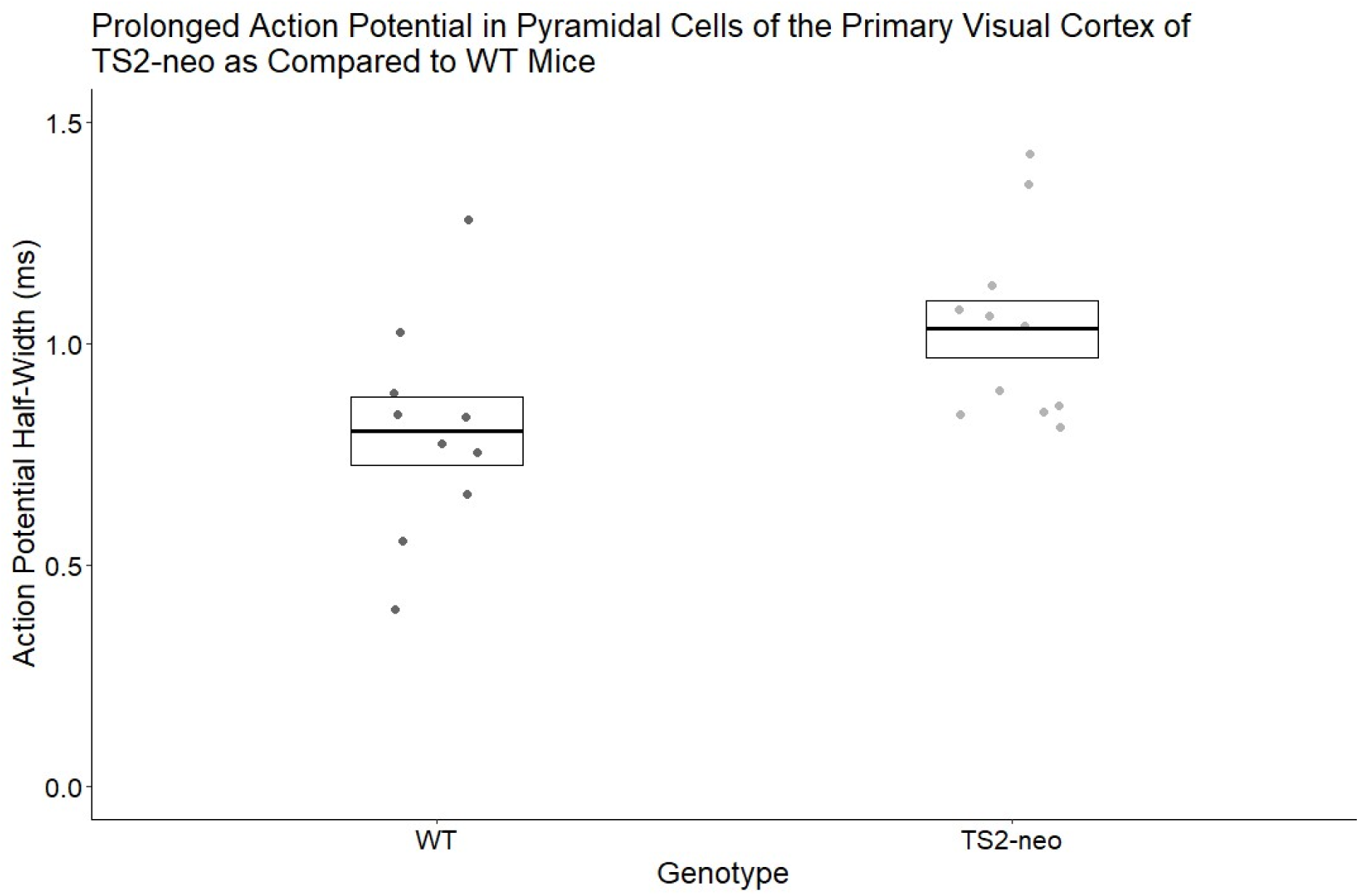
AP half-widths measured in V1 pyramidal cells of TS2-neo mice (right) and WT mice (left). Box plots show mean ± SEM, dots show values for each individual cell (WT: n=10, TS2-neo: n=11).

Further testing was conducted to explore whether the mutation had more of an effect on either the rising or falling phase of the AP waveform. This provided no evidence to suggest that TS2-neo heterozygosity impacts AP rise more than it does repolarisation of the membrane, or vice-versa. Genotype had a significant effect on the maximum rate of fall of the AP as well as on the total duration of AP rise and fall, while the effect on the maximum rate of rise of the AP was not quite significant. However, after multiple test correction none of these effects remained significant (maximum rate of rise [t-test]: t=1.7, P=0.051, corrected P= 0.31; maximum rate of fall [t-test]: t=2.2, P=0.023, corrected P=0.20; total rise time [t-test]: t=-2.0, P=0.032, corrected P=0.22; total fall time [t-test]: t=-2.1, P=0.025, corrected P=0.20). The values obtained for each of these properties from TS2-neo and WT mice is shown in Figure 3.

**Figure 3:**
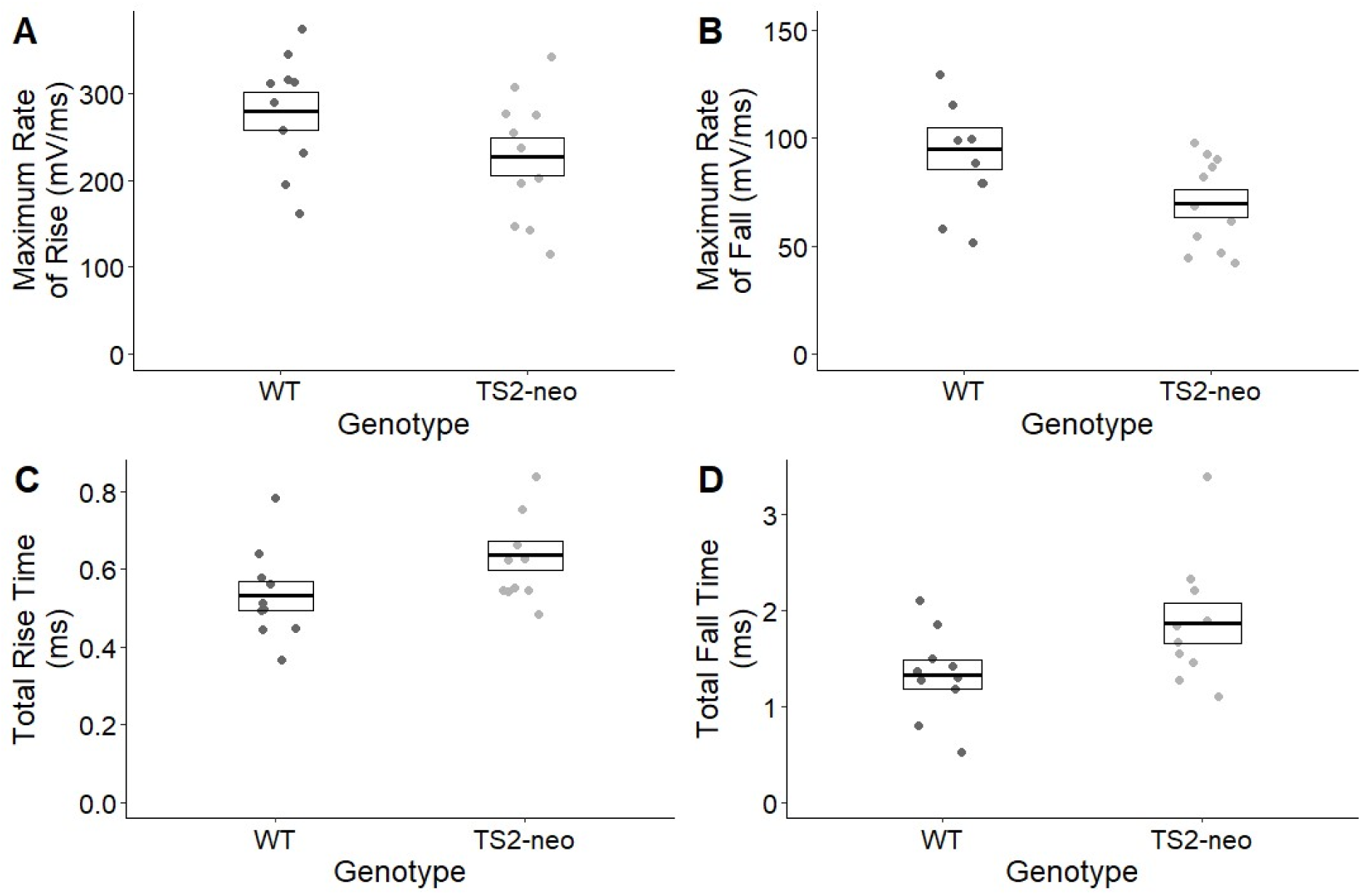
AP maximum rate of rise (A) and of fall (B), total rise time (C) and total fall time (D) compared for V1 pyramidal cells in TS2-neo mice and WT mice. Box plots show mean ± SEM, dots show values for each individual cell (WT: n=10, TS2-neo: n=11).

### Differences in neuronally measured contrast sensitivity functions (CSFs) from TS2-neo and WT mice

In order to assess the effect of the TS2 mutation on basic visual stimulus processing we measured CSFs of V1b neurons using two-photon calcium imaging. Visually responsive cells were detected as described in the Supplementary Methods section (see also Supplementary Figure 2, showing visually responsive GCaMP6f-labelled neurons responding to a stimulus of low SF [0.014cpd] and high contrast [100%]). Fluorescence traces from each visually responsive cell were processed and used to measure the cell’s responsiveness to visual stimuli of each SF tested and at each contrast level (see Supplementary Figure 3). For each neuron, a CSF was generated, showing how contrast sensitivity varies with SF of the gratings (see Supplementary Figure 4). CSFs for all cells of a WT and TS2-neo animal are shown in Supplementary Figures 5 and 6, respectively. Data from all visually responsive neurons of all animals from each genotype were used to generate a population CSF for WT and for TS2-neo animals as shown in Figure 4, showing the measured contrast sensitivities of all cells from animals of each genotype for visual stimuli of each SF tested. Data were analysed such that between-animal variance was accounted for. A dot plot showing the distribution of cells falling into each contrast sensitivity category for visual stimuli of each SF tested is shown in Figure 5.

**Figure 4:**
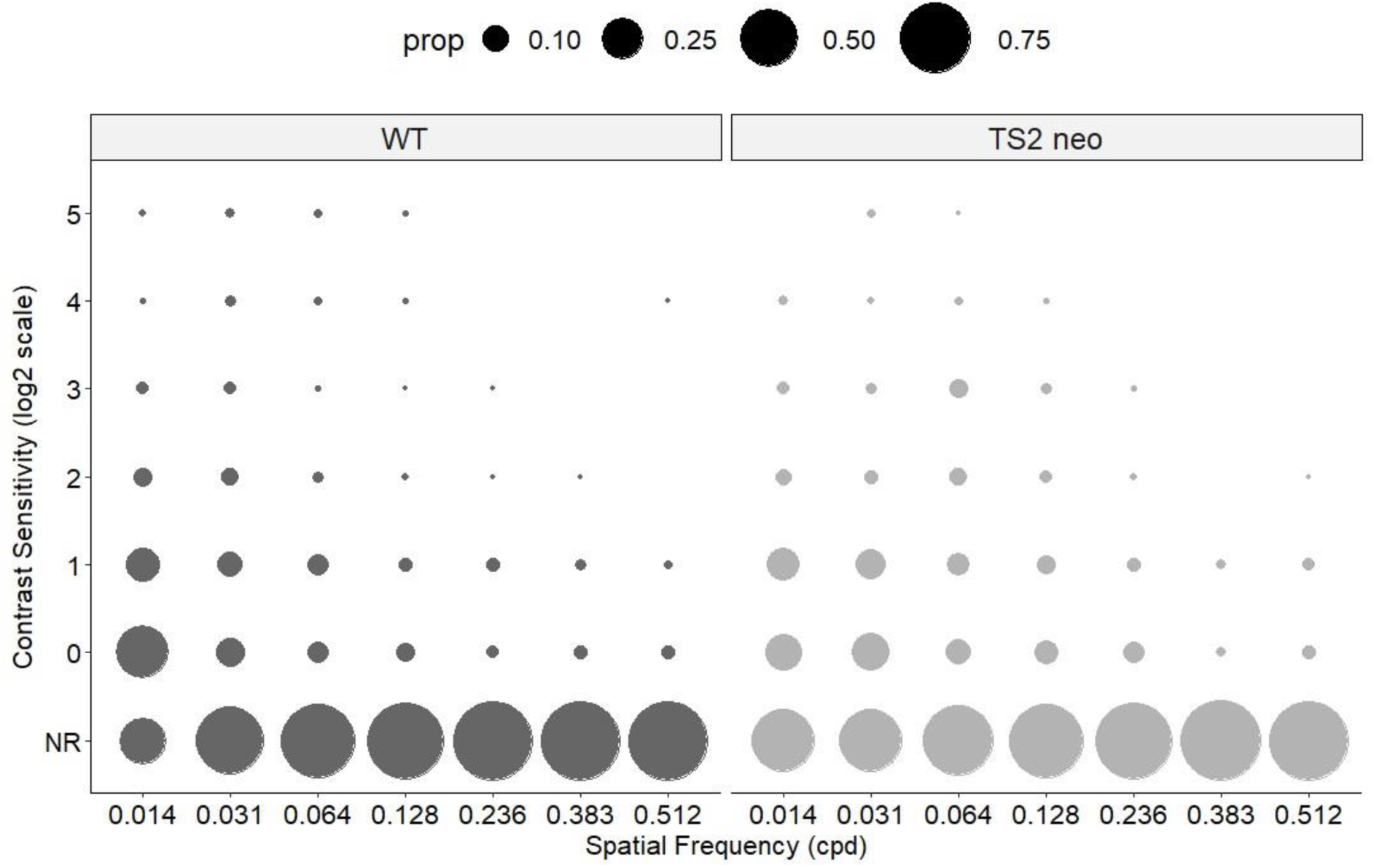
Contrast sensitivity of individual V1 neurons in all WT mice (left) and TS2-neo mice (right). Circle sizes represent proportions of cells responding at the contrast sensitivities given on the ordinate for each of the spatial frequencies shown on the abscissa, where 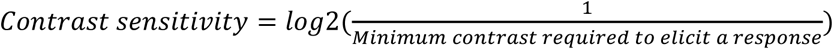. Circle sizes for proportions of 25%, 50% and 75%, respectively, are shown at the top. Note that there are proportionally more cells responding at lower spatial frequencies in WT mice and more cells responding at medium spatial frequencies in TS2-neo mice.

**Figure 5:**
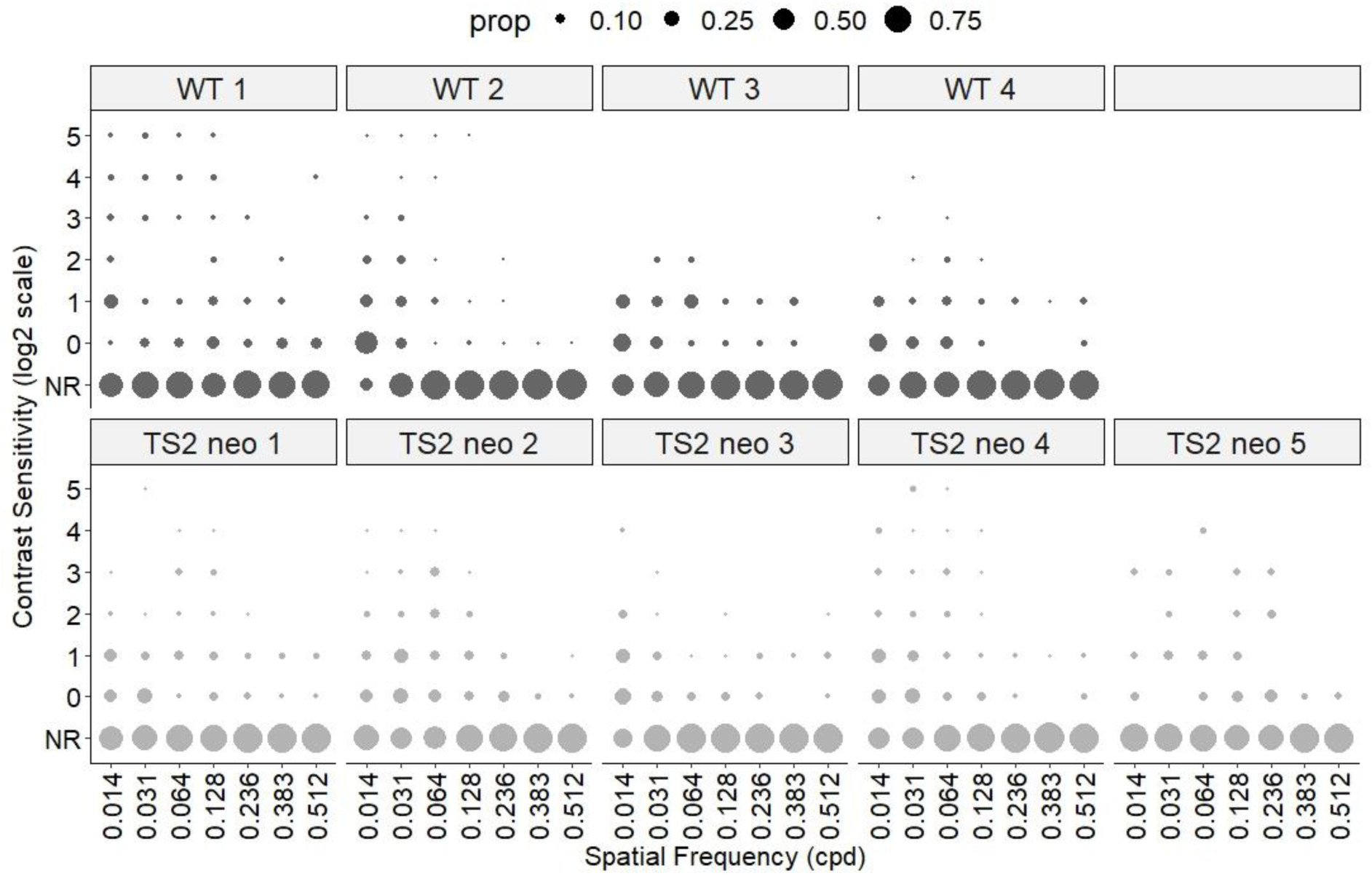
Contrast sensitivity of individual V1 neurons for 4 WT mice (top row) and 5 TS2-neo mice (bottom row). Format as in Figure 4.

A cumulative link mixed model was designed to assess how contrast sensitivity-variance-by-SF (an overall CSF from neurons recorded from) varies with genotype when between-animal variance is specified as a random factor. The assumptions of the model were deemed to be met based on visual inspection of residual distribution. The model was found to explain 17% of variance in contrast sensitivity measured from all cells (Cox and Snell [ML]: pseudo R^2^=0.17). This value is low as expected: the SF to which a V1b neuron responds varies based on the cell’s location (Zhang et al., 2015), cells recorded from in this experiment series were sampled across L2/3 V1b. Model testing found that the genotype-SF interaction term varied significantly with contrast sensitivity (Analysis of deviance: χ^2^=85, P=4.1×10^-16^, corrected P=1.2×10^-15^). This indicates that TS2-neo heterozygosity significantly affects the neuronally-measured CSF of the mouse.

Post-hoc testing was conducted to explore exactly how TS2-neo heterozygosity impacts neuronally measured contrast sensitivity to visual stimuli of both low (0.014cpd) and high (0.128cpd) SF. TS2-neo heterozygosity was associated with a reduction in contrast sensitivity to visual stimuli of low SF (0.014cpd [estimated marginal means]: Z ratio= −4.2, P=2.2×10^-5^, corrected P=2.4×10^-3^) and an increase in contrast sensitivity to visual stimuli of high SF, although significance was lost after correction for multiple testing (0.128cpd [estimated marginal means]: Z ratio= 2.73, P=0.0063, corrected P=0.063). This is mirrored in the proportion of all visually responsive V1b cells from TS2-neo as compared to those from WT animals which respond to high and low SF stimuli (Figure 6). 67.3% of neurons in WT mice respond at 0.014 cpd vs 41.4% in TS2-neo mice, while conversely only 9.4% neurons from WT mice respond at 0.128 cpd but 17.4% neurons from TS2-neo mice do.

**Figure 6:**
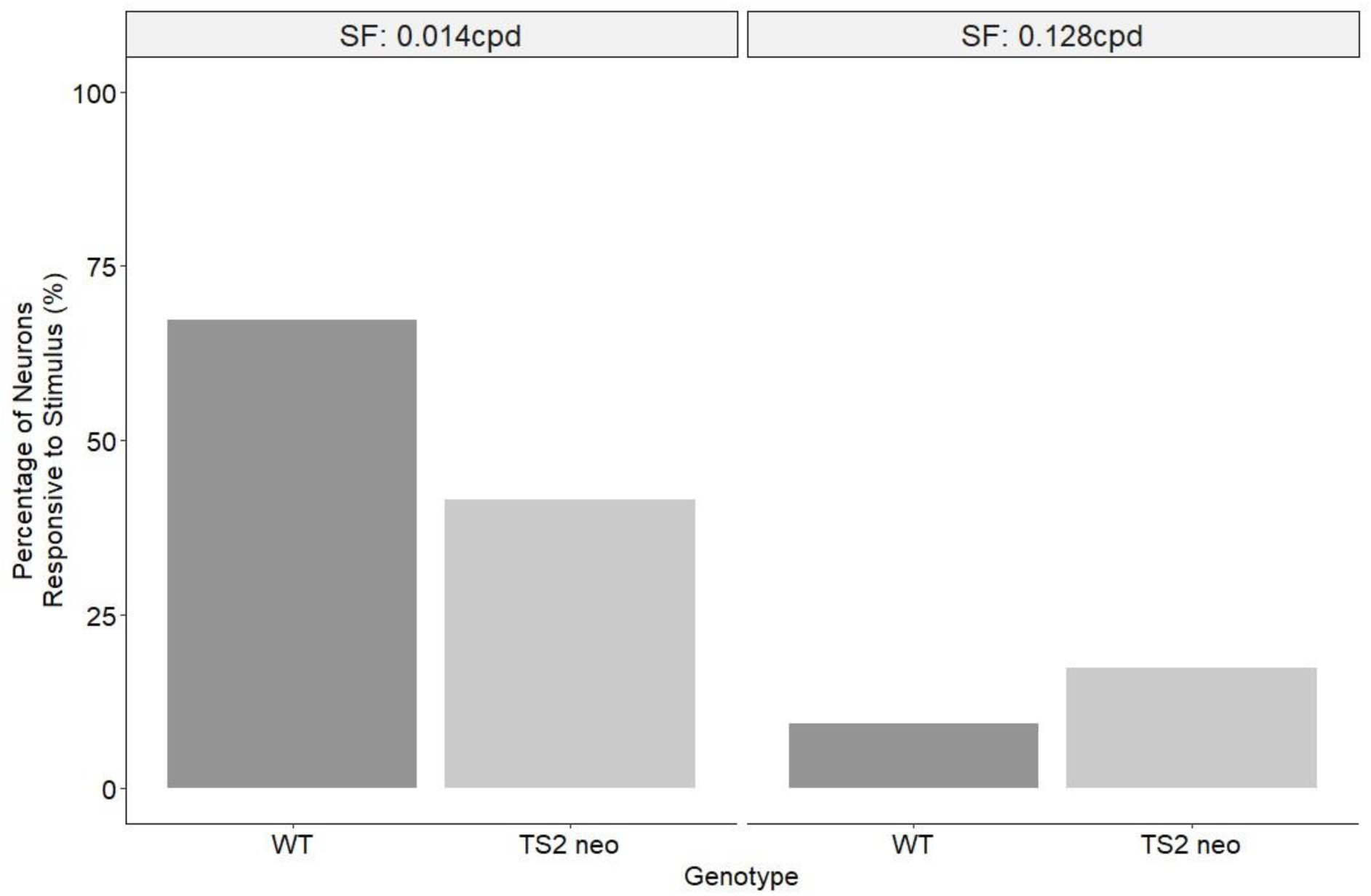
Proportions of V1b neurons from WT and TS2-neo mice responding to gratings of low spatial frequency (0.014 cpd, left) and proportions of V1b neurons from WT and TS2-neo mice responding to gratings of medium spatial frequency (0.128 cpd, right).

Overall, mice heterozygous for the TS2-neo mutation have significantly different neuronally measured CSFs as compared to WT mice. Neuronally measured contrast sensitivity to visual stimuli of low SF (0.014cpd) is significantly lower in mice heterozygous for the TS2-neo mutation as compared to WT. Neuronally measured contrast sensitivity to visual stimuli of high SF (0.128cpd) tends to be higher in mice heterozygous for the TS2-neo mutation as compared to WT.

### Increased PV+ cell density in V1 of TS2-neo as compared to WT mice

In order to ascertain whether the TS2 mutation affects the development of inhibitory interneurons in V1 we studied the distribution and density of PV+ cells, the most common GABAergic cell class in V1. Supplementary Figure 7 shows AlexaFluor488-labelled PV+ cells in sections from a TS2-neo and a WT mouse. PV+ cell density for sections collected from TS2-neo and WT mice was found to be normally distributed on visual inspection. A linear mixed regression method was used to model how PV+ cell density varied with genotype when controlling for between-animal differences. Model assumptions were found to be adequately met. Genotype was found to significantly impact V1 PV+ cell density for the model (analysis of variance: χ^2^=7.2, P=0.0075, corrected P=0.022). PV+ cell density measurements obtained from sections of WT and TS2-neo mice are shown in Figure 7. The model was found to describe 15% of the variance in the data, genotype was found to describe 9.2% of variance in PV+ cell density within V1 (conditional R^2^=0.15, marginal R^2^=0.092). The model predicted TS2-neo mice to have a 24% higher PV+ cell density in V1 as compared to WT (PV+ cell density in V1: WT= 92±7.7cells/mm^2^ [n=133 sections, N=4] mice; TS2-neo= 114±4.7cells/mm^2^ [n=198 sections, N=7 mice]).

**Figure 7:**
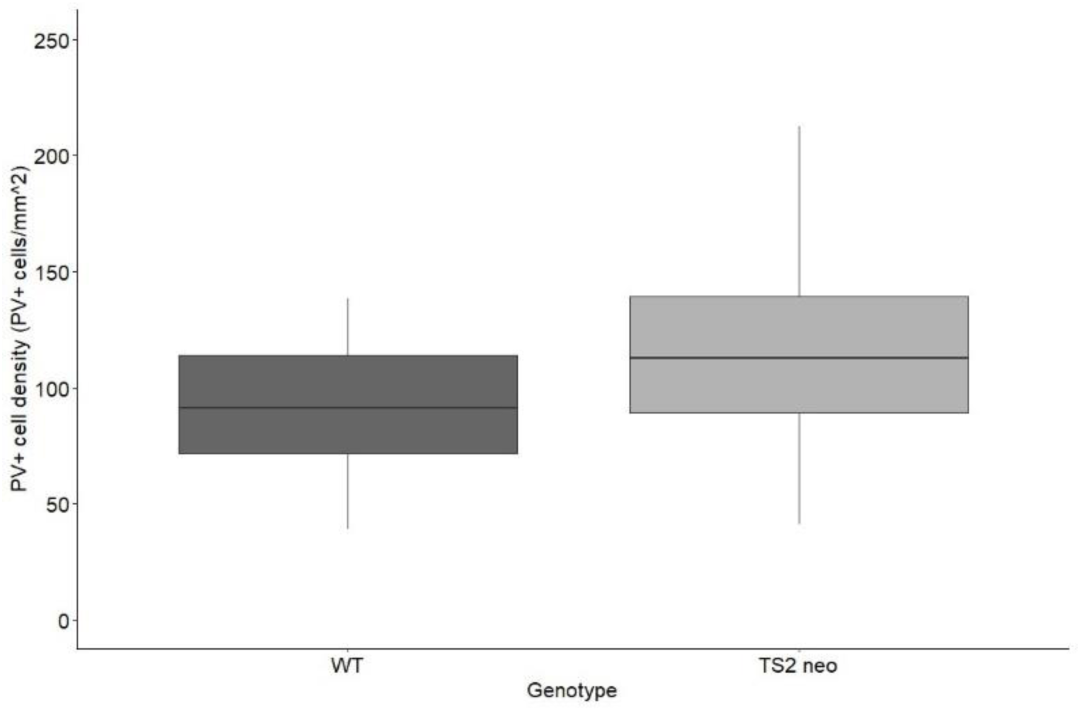
PV+ cell density in V1 of WT mice (left) and TS2-neo mice (right).The boxplot shows mean +/− IQR, The upper whisker (vertical line) extends from the hinge to the largest value no further than 1.5 * IQR from the hinge (where IQR is the inter-quartile range, or distance between the first and third quartiles). The lower whisker extends from the hinge to the smallest value at most 1.5 * IQR of the hinge. Data beyond the end of the whiskers are deemed outlying points and are plotted individually.

## Discussion

We found that the TS2-neo mutation prolongs action potentials in mouse V1 pyramidal cells as the TS2 mutation does in cardiomyocytes. It alters basic visual processing in V1b of mouse at the neuronal level: the CSF is characterised by decreased sensitivity to low SF and increased sensitivity to medium-to-high SF. Moreover, PV+ cell density is increased in V1.

### Prolonged pyramidal cell AP

Our results whole-cell patch clamp recordings in acute ex-vivo slices from TS2-neo mouse brains replicate those of (Paşca *et al*. 2011) in human iPSC-derived neurons. They provide evidence that classical TS mutations result in a prolonged AP in neurons as they do in cardiac myocytes (Splawski *et al*. 2004; Splawski *et al*. 2005; Dick *et al*. 2016). Our data do not give an indication by which mechanism neuronal AP is prolonged since rise times and rates were affected similarly as fall times and rates. We found no difference in spike frequency adaptation in TS2-neo mouse pyramidal cells as compared to WT.

The TS2 mutation has recently been found to prolong the AP duration in adrenal chromaffin cells (Calorio et al. 2019). Chromaffin cells have APs which are largely underpinned by sodium and potassium currents, similar to those which govern APs in neurons. In the chromaffin cell, the TS2 mutation causes a reduction in NaV1 channel density, which slows the rate of depolarisation of the AP, resulting in a prolonged AP (Calorio *et al*. 2019). CaV1.2 is known to impact gene transcription and ion channel trafficking within the neuronal membrane (Tian et al. 2014; Servili et al. 2020).

It is therefore more likely that the prolonged neuronal AP is caused by one or both of either a direct result of prolonged depolarising currents through the mutant CaV1.2 channel, or by a mutation-related impact on NaV1 channel density in the neuronal membrane.

### Abnormal contrast sensitivity function

Neurons from TS2-neo mice were less likely to respond to visual stimuli of low SF, but more likely to respond to visual stimuli of mid-high SF compared to WT mice. These results point to a basic visual processing abnormality in TS2-neo mice.

Three TS2 patients reported in the literature have been found to have abnormalities in eye movement and function, with comorbid neurological disorder (Splawski *et al*. 2005; Walsh *et al*. 2018). While abnormalities in visual perception have not been reported in TS2 patients, our finding of abnormal visual function in the TS2 neo mouse suggests that the TS2 mutation, and perhaps TS mutations in general may be associated with abnormalities in visual processing. This hypothesis is supported by a case report from a patient with atypical Timothy Syndrome (TSA) who exhibited severe visual dysfunction (Gillis et al. 2012).

It is possible that TS mutations result more generally in abnormalities in sensory perception. A study by (Rendall *et al*. 2017) found that TS2-neo mice had a superior ability to respond to silent gaps within auditory stimuli compared to WT mice. It should be noted that TS is commonly associated with ASD: around 80% of TS patients meet the diagnostic criteria for ASD (Splawski *et al*. 2004). ASD often presents with abnormal sensory processing symptoms across all sensory modalities (Marco *et al*. 2011; Robertson and Baron-Cohen 2017). ASD patients have been found to be superior to neurotypical subjects in processing high spatial frequency (SF) visual stimuli (Vlamings *et al*. 2010; Kéïta *et al*. 2014). (Kéïta *et al*. 2014) found that ASD patients have an enhanced contrast sensitivity for simple visual gratings stimuli of high SF.

As such, the TS2-neo mouse model of TS has also been considered as a mouse model of ASD of a specific known genetic cause (Bader et al. 2011); (Bett *et al*. 2012); (Rendall *et al*. 2017). The superior auditory processing ability in TS2-neo mice found by (Rendall *et al*. 2017) is similar to those previously reported in ASD (Bertone *et al*. 2005) (Mottron *et al*. 2006). (Cheng *et al*. 2020) found that the inbred BTBR mouse model of idiopathic ASD had higher contrast sensitivity to mid-high SF visual stimuli as compared to WT mice. In this study, we found that the TS2-neo mouse model had increased contrast sensitivity to visual stimuli of mid-to-high SFs but not at the highest SF tested (0.512cpd). It is not clear yet how increased AP width might lead to an increase in sensitivity to visual stimuli of higher spatial frequency, but it is feasible that wider APs affect the temporal integration of bursts of action potentials elicited by grating stimuli, resulting in a stronger response to stimuli that in WT animals remain subthreshold.

Overall, our finding that TS2-neo mice has an abnormal CSF was similar to that found in ASD patients by (Kéïta *et al*. 2014) and in the BTBR mouse model of ASD by (Cheng *et al*. 2020). This suggests that the TS2-neo mouse has basic visual processing abnormalities which resemble those seen in ASD.

### Increased PV+ cell density

Inhibitory interneuron dysfunction is thought to be involved in the pathology of ASD (Hussman 2001) and epilepsy (Magloire et al. 2019), two common neurological phenotypes of TS. PV+ cell number has been found to be abnormal in various areas of the brain in people with ASD by postmortem histological study (Lawrence et al. 2010; Zikopoulos and Barbas 2013; Ariza et al. 2016; Hashemi et al. 2017). Moreover, the gene coding for PV is the most strongly downregulated in the ASD brain as compared to neurotypical subjects by postmortem transcriptome (Schwede et al. 2018). This evidence has led to the PV Hypothesis of ASD (Filice et al. 2020). A similar theory suggests that abnormal PV+ cell activity may result in epileptiform neural activity in the brain (Magloire *et al*. 2019). The evidence from this study supports the hypothesis that the ASD-related classical TS2 mutation results in abnormalities in PV+ cell density in the adult mouse brain.

Abnormalities in interneuron migration have been demonstrated in the developing brain of TS2-neo mice (Horigane *et al*. 2020). Beyond that, TS-related abnormalities in the development and density of PV+ interneurons have not previously been explored in animal models. However, PV+ cell number abnormalities have been reported for various brain areas in mouse models of ASD (Tripathi et al. 2009; Allegra et al. 2014; Lauber et al. 2016; Paterno et al. 2021; Yang et al. 2021; Kourdougli et al. 2023). Specifically, PV+ cell number has been found to be increased by 20% in V1 in the En2 KO mouse model of ASD as compared to WT (Allegra *et al*. 2014); the En2 gene encodes the homeobox protein engrailed-2, which is involved in embryonic development and is implicated in ASD (Benayed et al. 2009). Our study supports this finding; however, the sample size was relatively small, and further studies are needed to conclusively demonstrate an effect of classical TS mutations on PV+ cell development, number and function in the mouse brain.

How might an increase in PV+ cell density affect neuronal CSFs? Visual gain control is modulated by different classes of GABAergic interneurons, including PV+ cells, somatostatin-positive and vasopressin-positive cells (Pakan et al. 2016). A higher density of PV+ cells might change the local balance of excitation and inhibition. Increased surround inhibition could result in a smaller excitatory receptive field centre and therefore higher contrast sensitivity at higher spatial frequencies (Sengpiel et al. 1997).

In summary, our results show that classical TS mutations a) cause abnormal electrical activity and properties in single neurons in the mouse brain; b) PV+ interneuron development and distribution in the brain; and c) result in abnormalities in basic sensory processing within the visual domain. Study of the TS2-neo mouse model of TS is useful for gaining a better understanding of the neurophysiology of TS.

## CRediT authorship contribution statement

Rosie Craddock: Methodology, Investigation, Software, Formal analysis, Writing – original draft, Writing – review & editing, Visualization; Cezar Tigaret: Conceptualization, Funding acquisition, Methodology, Supervision, Writing – review & editing; Frank Sengpiel: Conceptualization, Funding acquisition, Project administration, Supervision, Writing – original draft, Writing – review & editing, Visualization

## Acknowledgements

This study was supported by a PhD studentship to RC under the SWBio Doctoral Training Programme funded by the Biotechnology and Biological Sciences Research Council and a MRC New Investigator Research Grant (MR/V034111/1) awarded to CMT. We thank Fangli Chen for expert technical assistance.

