## Supplementary Material for "Disruptions in Primary Visual Cortex Physiology and Function in a Mouse Model of Timothy Syndrome"

#### Supplementary Methods

*Passive membrane properties*

The voltage trace corresponding to the -50pA current step was used to measure passive membrane properties of the cell as well as voltage rebound and voltage sag. The passive membrane properties measured were membrane input resistance, capacitance, and the membrane time constant. The methods used to obtain these values were based on those reported by (Tamagnini et al. 2015). An illustration of how these measurements were obtained are shown in Supplementary Figure 1. Resting membrane potential was taken as the membrane potential of the cell measured 100ms before the onset of the hyperpolarising current. This was typically -70mV, and was not based on empirical measurement.

Rheobase was measured empirically as the minimum input current required to evoke an AP for the cell, measured in pA. No mathematical calculation was used to estimate rheobase to a higher degree of accuracy. The maximum instantaneous firing frequency of each cell was measured as the reciprocal of the minimum measured time between any two spikes for any input current to the cell. This was measured in Hz. Minimum AP onset latency was measured as the minimum time between stimulus onset and AP firing (taken from AP threshold). Minimum AP onset latency was measured in ms.

*Detection of cells in two-photon imaging*

Cell fluorescence traces were smoothed using a 100-point moving window and were down sampled 5 times. The mean fluorescence of the cell at the baseline period for a given stimulus was compared to that for the period during which each stimulus was shown. Cells which had a significantly higher fluorescence during the stimulus presentation as compared to the baseline period for stimuli of SFs shown at either 100% or 50% contrast were deemed to be visually responsive. The criteria were that the fluorescence had to be significantly higher during the stimulus as compared to the baseline period by Wilcoxon Signed Rank testing (threshold value = 0.01) with the average fluorescence being 30% higher for the stimulus as compared to the baseline period. Cells deemed to be visually responsive (to stimuli of any SF) were further tested to find which SFs they responded to, and the minimum contrast at which they responded to for that SF. Again, this involved Wilcoxon Signed Rank testing (threshold value = 0.01) and a 30% increase in fluorescence between the stimulus presentation and baseline periods. Data processing codes were written in MATLAB 2022a and are available on GitHub.

*Contrast response functions of visually responsive neurons*

Contrast sensitivity was classified in one of 7 categories for each of the 7 SFs (ordered from high to low), with both the independent (SF) and dependent (contrast sensitivity) variables being ordinal. A form of contrast sensitivity function (CSF) for cells from mice of each genotype was created by measuring the number of cells from mice of each genotype falling into each sensitivity category for stimuli of each SF. To assess how CSF varies by genotype, a cumulative link mixed model regression was completed where between-animal variance was taken as a random factor. The assumptions of the model were tested graphically and were found to be met. The goodness of fit of the model was tested using Cox and Snell method included in the nagelkerke function of the rcompanion package (Mangiafico 2023).

*Statistical analysis of PV+ cell density*

Analysis was completed in R. PV+ cell density data were visualised to assess distribution. A linear mixed model was used to explore how genotype impacted PV+ cell density in V1 while controlling for between-animal variance. This involved use of the lme4 package in R (Bates et al. 2015). Homoscedasticity was assessed visually by assessing equality of residuals across the fitted values of the model by eye, while normal distribution of the model residuals was tested using the Shapiro-Wilk method. The model was deemed to meet assumptions of homoscedasticity (not shown) and of normally distributed residuals (Shapiro-Wilk: W=0.99, P=0.23). The goodness of fit of the model was tested by estimating pseudo $R^2$ values which were obtained using the MuMIn package in R (Barton 2024) which uses methods based on those of (Nakagawa and Schielzeth 2013). The effect of genotype on PV+ density in V1 was assessed by comparing the statistical model described above against a null model—which accounted for between-animal variance but did not model PV+ cell density by genotype—using analysis of variance. Effect size was estimated from the model directly.

### Supplementary figures

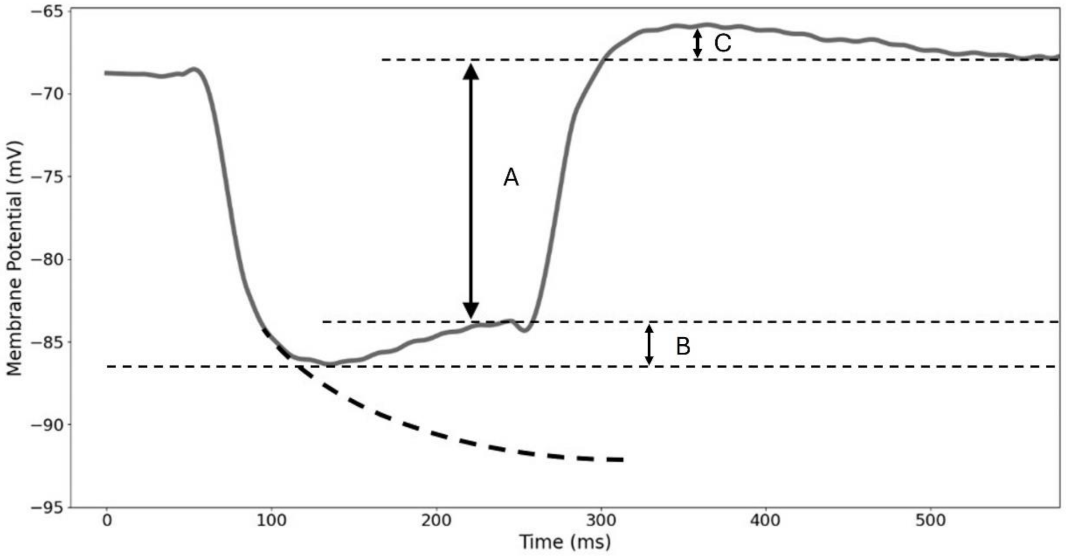


**Supplementary Figure 1**: Quantitative measurement of AP height from patch-clamp recordings. Action potential threshold was measured as the membrane potential at which the rate of change of membrane potential reached 15.2mV/ms (as per (Paşca et al. 2011). The membrane potential half-way between the threshold potential and the full AP amplitude (A) was designated the AP half-height. The voltage difference, A, divided by the input current (-50pA) is taken as the input resistance (MΩ). The measurement B was used to give the voltage sag (mV). The measurement C was used to obtain the membrane potential rebound (mV). An exponential fit of the curve of the depolarisation (based on methods of Tamagnini et al. (2015), indicated by the dashed line) was used to obtain a measure of the membrane time constant, τ (ms). Membrane capacitance was taken as τ/input resistance (pF).

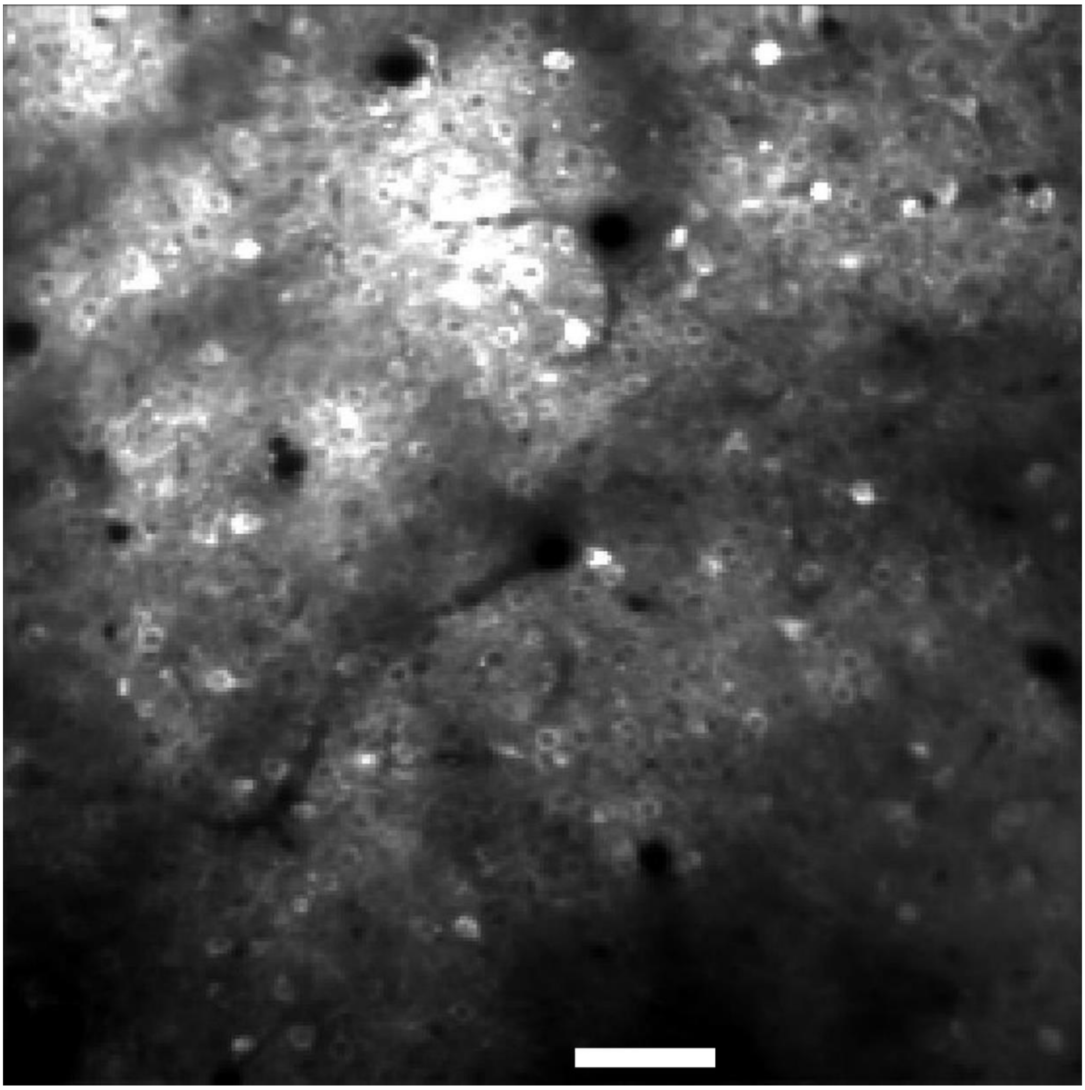

**Supplementary Figure 2**: GCaMP6f-labelled neurons in V1 of a TS2-neo mouse responding to a grating stimulus of low SF (0.014 cpd) and high contrast (100%).

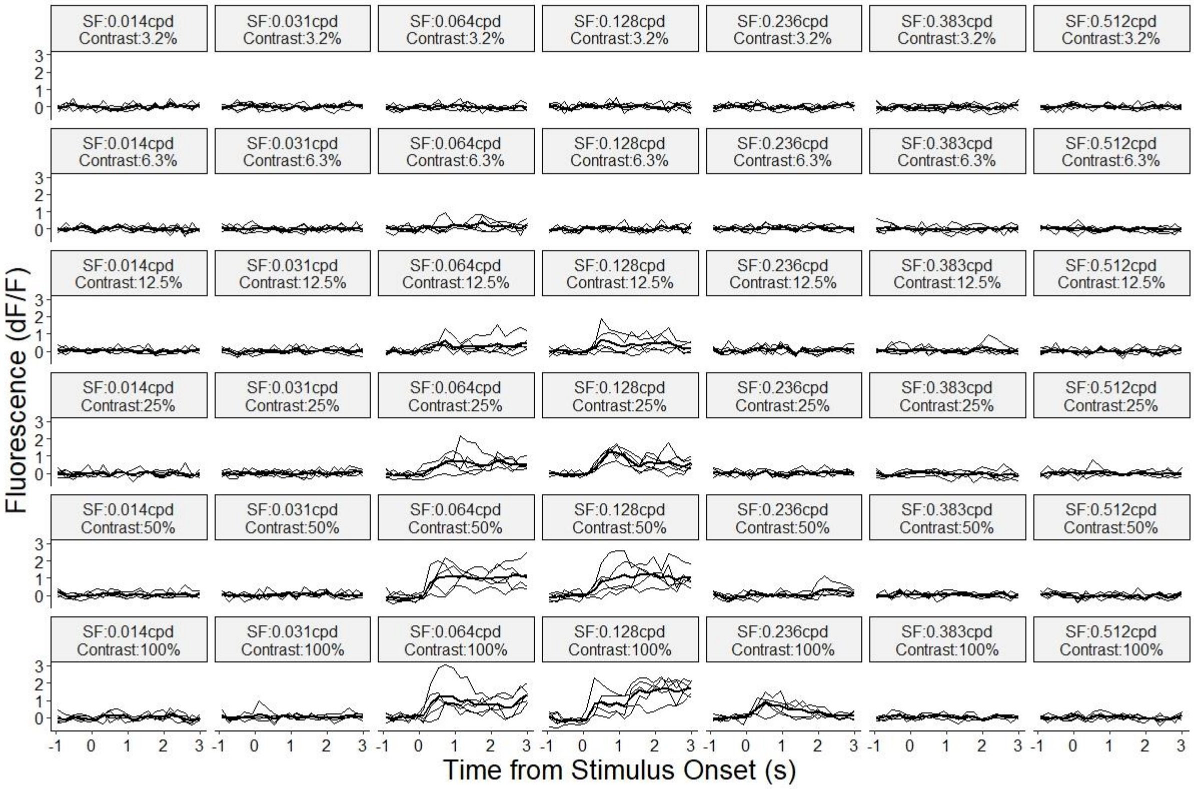


**Supplementary Figure 3**: Fluorescence traces from a visually responsive V1 neuron in response to grating stimuli of 7 different SFs  and 6 contrast levels tested.

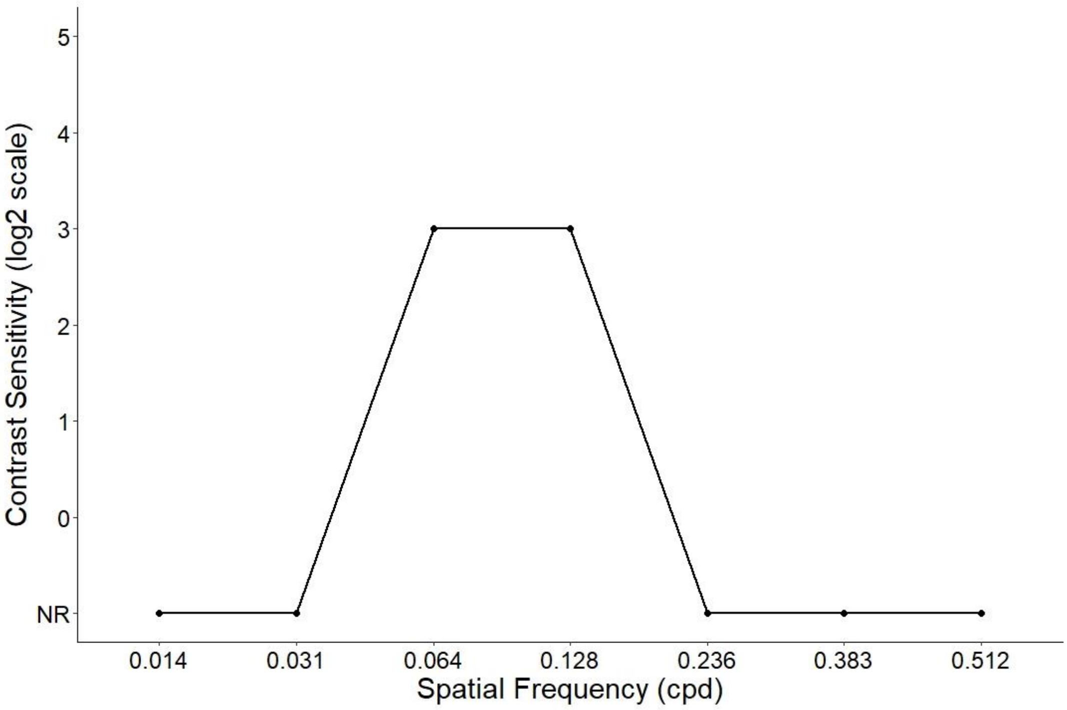


**Supplementary Figure 4**: Example contrast sensitivity function of an individual V1 neuron (the cell shown in Supplementary Figure 3).

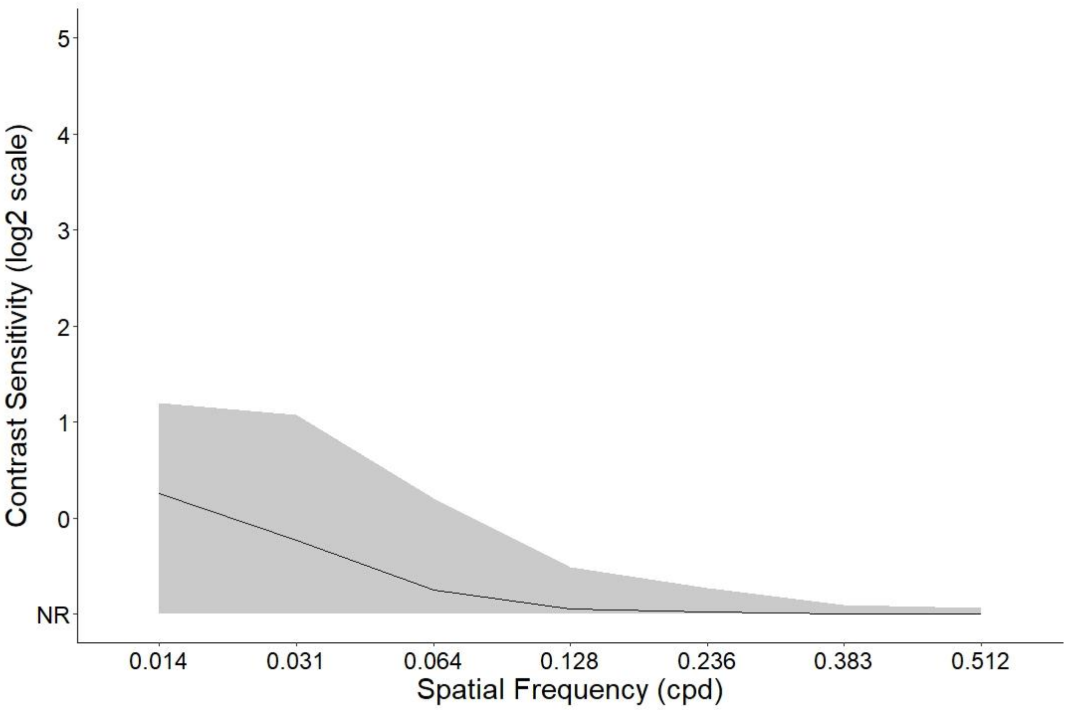


**Supplementary Figure 5**: Population CSF for one WT animal. The black line represents the mean, the grey ribbon represent the inter-quartile range (IQR).

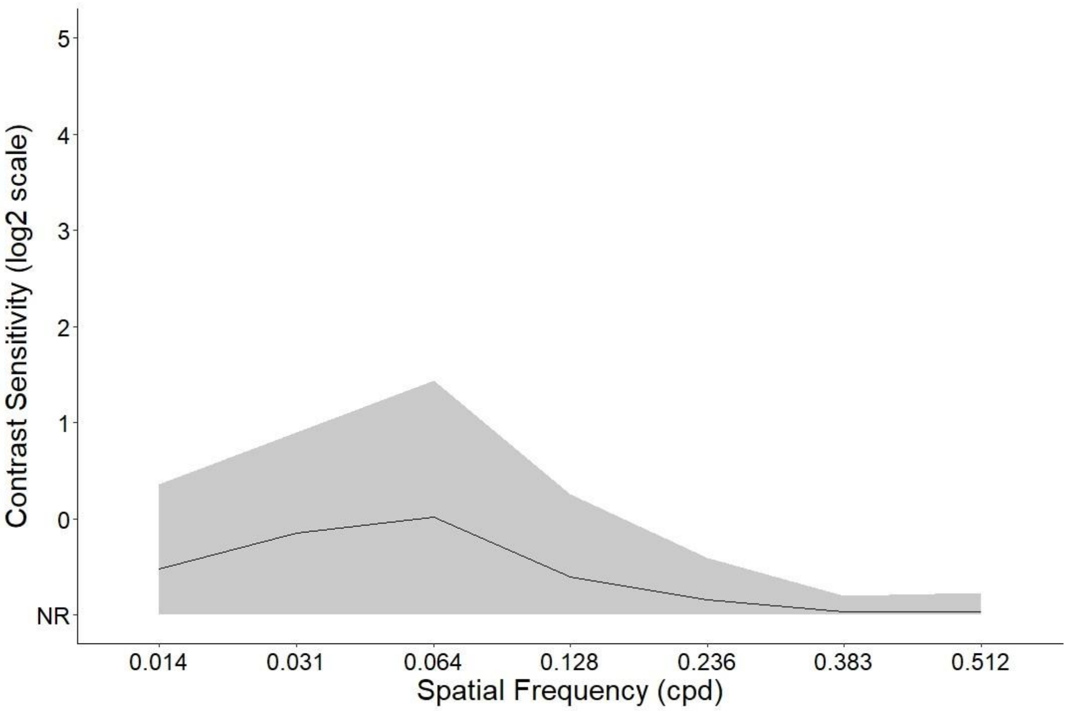


**Supplementary Figure 6**: Population CSF for one TS2-neo animal. The black line represents the mean, the grey ribbon represent the IQR.

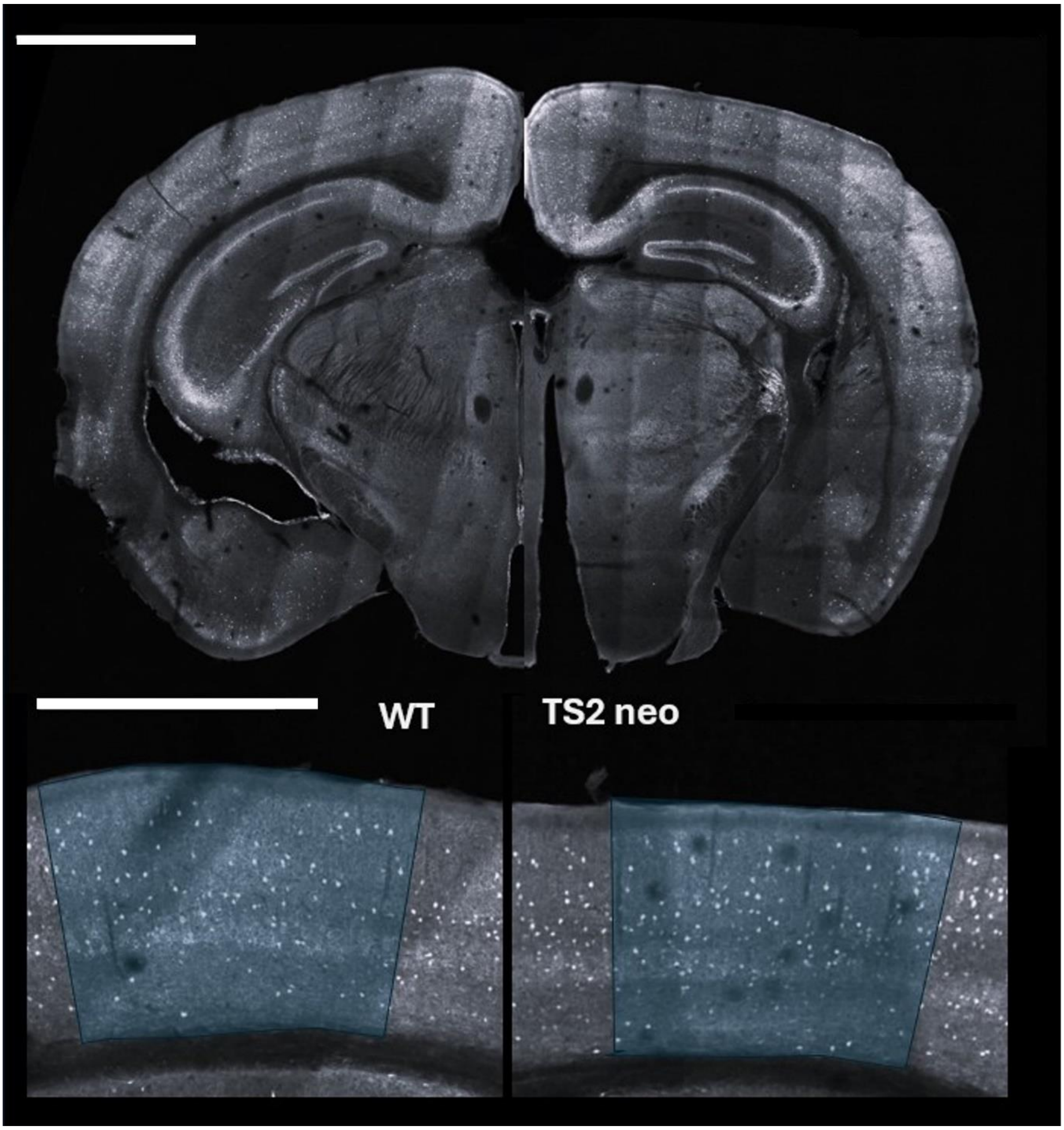


**Supplementary Figure 7**: AlexaFluor488-labelled PV+ cells in sections from a WT mouse (left) and TS2-neo moue (right). The bottom images show V1 in higher magnification (highlighted in blue). Scale bars, 2 mm for the top panels and 1 mm for the bottom panels.
